# Characterization of phage-antibiotic interaction through different *in vitro* methods: case study of a vibriophage

**DOI:** 10.64898/2026.05.26.727873

**Authors:** Brunel Archambeaud, Clément Douarre, Pierre R. Marcoux

## Abstract

Climate change and warmer oceans will amplify the impacts on public health of waterborne harmful microorganisms. Phagotherapy offers a promising alternative; but as of today, phages can only be administered to patients when delivered along with antibiotics. Understanding possible interactions between these agents – indifference, synergy or antagonism – is thus a pivotal point. While several methods exist for characterizing such interaction, consensus on a reference method is still lacking. In this work, we screen and compare several *in vitro* characterization methods, using as a model nt-1, a phage of *Vibrio natriegens*, and studying its interaction with cefotaxime, a 3G cephalosporine. The different methods highlight different aspects of the interaction, depending whether they focus on phage or bacterial biomass. Overall, we see evidence of antagonism between the studied phage and antibiotic: this antagonism is at its optimum for antibiotic concentration of minimum inhibitory concentration (MIC)/2. Given the non-linear nature of interaction, it appears essential to use multiplexed methods and to cross technics.

**AUTHOR SUMMARY:** Currently, antimicrobial resistance results in close to one million victims per year worldwide. In response to this alarming situation, new antimicrobial drugs and alternative therapies with innovative mechanisms have to be developed, such as phage therapy. It relies on the use of specific bacterial viruses, called bacteriophages (phages), that are therefore natural antibacterial agents. This therapy is strongly investigated for its potential to stop bacteria whenever antibiotics are no longer effective. Phage therapy is a highly personalized approach especially because of the narrow specificity of phages. Understanding how the efficiency of phages could be improved by the use of other antimicrobials, such as antibiotics, is essential in the fight against pathogens. Using a combination of a phage and an antibiotic, instead of only an antibiotic, imposes to think about new in-vitro tests for susceptibility testing. In the particular case of *Vibrio* bacteria, a common genus of waterborne pathogens, we investigated the efficiency of a phage in presence of cefotaxime, a last resort antibiotic, through different in-vitro methods, in liquid phase as well as on agar media. We observed a decreased efficiency of the phage, in other words an antagonism, especially at the lowest concentrations.

## INTRODUCTION

Among waterborne pathogens, Vibrios are a globally important source of bacterial diseases in human and aquatic animals. These Gram-negative bacteria are commonly found in warm, slightly salty waters and their growing prevalence in aquatic ecosystems mirrors the global warming(1–4). Several reports have recently indicated that human vibriosis are increasing worldwide, including lethal acute diarrheal diseases, such as wound infections, gastroenteritis and bloodstream infections(5,6). Furthermore, a high prevalence of *Vibrio* pathogenic strains has been recently reported by EFSA in seafood(7). Unfortunately, these serious opportunistic human pathogens are also affected by the phenomenon of antibioresistance(7–10). In this context, bacteriophages (phages), prokaryotic viruses first discovered one century ago on *Vibrio cholerae*(11), can infect and lyse bacteria. That is why they have raised renewed interest as alternative treatment(12,13) against antibiotic resistant strains. Since they are themselves subject to antimicrobial resistance, it is of great interest to evaluate their impact(14–18) when administered jointly with antibiotics, a field of study termed Phage-Antibiotic Interaction (PAI). In particular Phage-antibiotic synergy (PAS) is raising hopes of an enhanced treatment effect and is as of today one of the major research axes in the phagotherapy field(19,20) . However, combining phages and antibiotics is not always beneficial. Antagonism is a phenomenon much less studied but not infrequent(21). To prevent therapy failures, it is necessary to study this phenomenon as well as understanding its underlying mechanisms. The purpose of the present study is to investigate the PAI between a model of vibriophage, nt-1, and cefotaxime considered as a model of 3G-cephalosporin, a class of broad-spectrum β-lactam antibiotics frequently used as a last resort antimicrobial chemotherapy.

To characterize PAI, a range of *in vitro* methods exist, which differ regarding their interpretation of synergy. A first family of techniques, biomass-focused methods, defines synergy as the ability of two antimicrobial agents to have a higher impact when administered together, compared to the addition of their effects when delivered independently(22). Hence, the possible interaction effects are absence of effect, addition, synergy and antagonism. These methods include:

- time-kill curve by optical density (OD)(17,20,23),
- optical assessment of bacterial respiratory activity (24,25),
- imaging and counting of individual cells(15).

Biomass-focused methods are usually performed in broth. Tests can be multiplexed in multiwell plates to screen the effect of different antimicrobial agents at different concentration ranges. From these so-called checkerboard assays, several strategies exist to process the data and conclude on the interaction effect. Gu Liu *et al.*(26) described a model of kinetic curves and interactions plots for each of the possible interactions. In an attempt to synthesize interaction in a single value, many studies(27–30) calculate a Fractional Inhibitory Concentration Index (FICI)(22), based on the ratio of the minimum of inhibitory concentration of each antimicrobial agent. Based on this factor value, conclusions can be drawn regarding the type of interaction: synergy (<1), indifference (between 1 and 2) or antagonism (>2).

Different from these biomass-focused methods, some phage-focused methods have been developed. Among them, Comeau *et al.*(31) define PAS as an enhanced production of phages by the host submitted to a sublethal concentration of antibiotic. Methods focusing on this effect, that we will call phage-focused methods, include:

- lysis plaque morphology and size on Petri dish(31),
- one-step growth assay(31–33).

Finally, Mulla *et al.*’s approach(34) is a somewhat mixed method : they measure biomass and the delay after which bacterial population collapses, linking it to the phage production rate through a mathematical model they called PHORCE.

As we can see, there exists multiple methods to measure and assess PAS, and there is no single method used as a standard. In this context, how should we reliably assess the interaction of a given phage-antibiotic couple? In this study, we dissect and review the relative merits and limitations of different methods and the answers they provide, as well as the links that can be found between them. We use a model of Vibriophage, nt-1, whose host is *Vibrio natriegens*. This bacterium is of high interest as a model as it is in the same genus as pathogenic bacteria such as *V. haemolyticus* or *V. vulnificus*. We study the virulence of nt-1 phage – a phage discovered in 1976(35) but understudied since then – in the absence and presence of sublethal concentration of cefotaxime, a 3G cephalosporin that has shown great synergy potential with *E. coli* phages^4^. Indeed, like many cephalosporins, at sublethal concentration, cefotaxime affects the cell division, preventing the daughter cells to fully separate, causing the bacteria to grow filamentous. Since their surface area is enlarged, filamentous bacteria make easier targets for phages since they are presumed to show larger adsorption surfaces, which facilitates infection by phages. Furthermore, by slowing down the holin accumulation in the membrane, cefotaxime causes lysis to be delayed, leading to an increased phage production, since they have more time to mature inside the infected cell(36). Filamentation is thereby reputed a key factor of phage-antibiotic synergy(37).

In this work, we study interaction between cefotaxime and nt-1 phage through both biomass-focused methods and phage-focused methods. As a first experiment, we observed lysis plaque size and morphology on agar in presence and absence of antibiotic. To be able to test more precisely combination of a range of antibiotic and phage concentrations, we perform checkerboard assays with both measurements of OD over time and determination of bacteria survival at a final point via a colorimetric assay of respiratory activity. We can thus observe an antagonism at sublethal concentrations of the antibiotic. To better understand this antagonism, we determine effect of cefotaxime at 0.17mg/L, the sublethal concentration that shows highest antagonistic effect while considering bacteria survival, on phage virulence factors, burst size and latency time.

## RESULTS

### Lysis plaques morphology and size on a Petri dish

According to Comeau *et al.*(31), PAS can be demonstrated by placing an antibiotic diffusion disk on an agar plate seeded with both the host bacteria and the studied phage. Comeau *et al.* obtained spectacular results especially with cephalosporins such as cefotaxime on *E. coli* MFP: lysis plaques of phiMFP close to the antibiotic inhibition disk were much larger than in the rest of the plate. When following the same protocol for the *V. natriegens* / nt-1 / cefotaxime triad, a similar effect on lysis plaque size is observable, although the size differences were not as dramatic as in Comeau *et al.*’s experiments (Figure 1). Indeed, at the limit in which effect of the antibiotic is no longer observable on the bacterial density, lysis plaques are bigger than the one in the outer zone considered as control. Moreover, we note another more surprising effect in the area where bacterial density is lower but growth is not fully inhibited: lysis plaques are smaller than controls.

**Figure 1.**
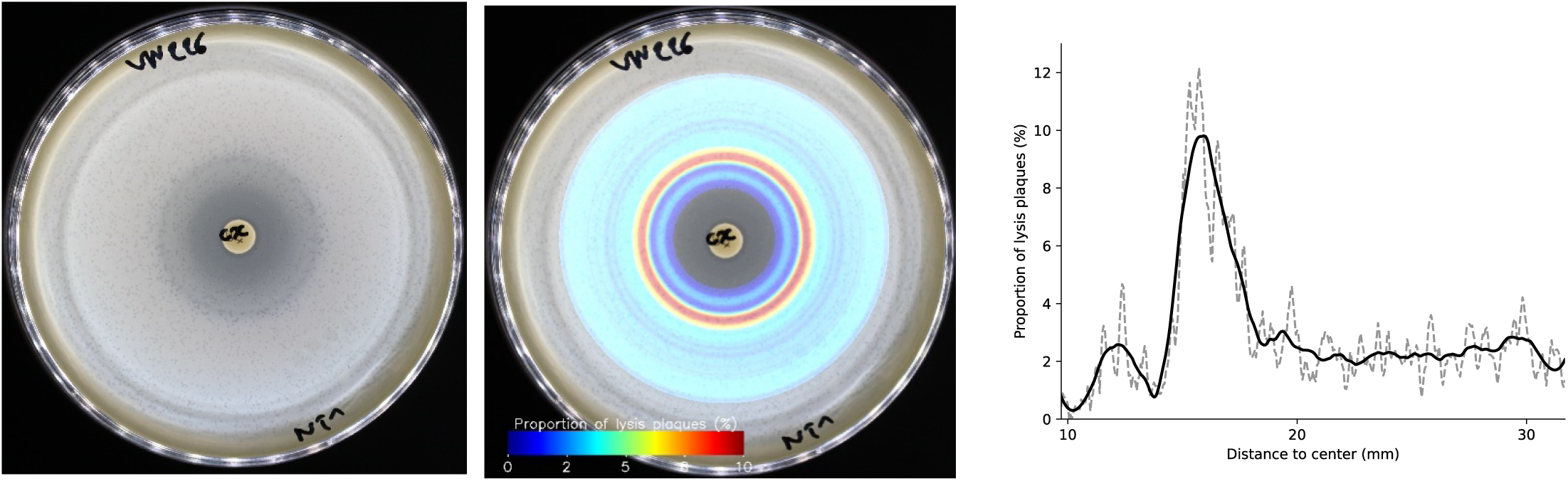
: Lysis plaque of nt-1 phage in presence of antibiotic disk. Left: Picture of Petri dish containing an antibiotic disk inoculated with phage, 24 hours after inoculation. Middle: lysis plaque density is measured in concentric circles and translated in color gradient, from red to indigo: red is higher plaque density, indigo is lowest. Cyan zone is considered as control where AB has no more effect. Right: plot of the lysis plaques density vs distance to inhibition disk.

### Combining phage and antibiotic in agar

Following Comeau *et al.,*(31) we evaluated PAI on agar enriched with phage and subinhibitory concentration of antibiotic. A control dish was seeded in parallel with the same amount of phages and bacteria as in the PAI settings, but without any other repressor.

The morphology of the plaques is indeed very different in absence and presence of antibiotic, but this difference is of another nature than the one displayed in Comeau *et al.* experiments. In our experiments, plaques appeared much smaller and with a turbid aspect in the presence of cefotaxime (Figure 2).

**Figure 2.**
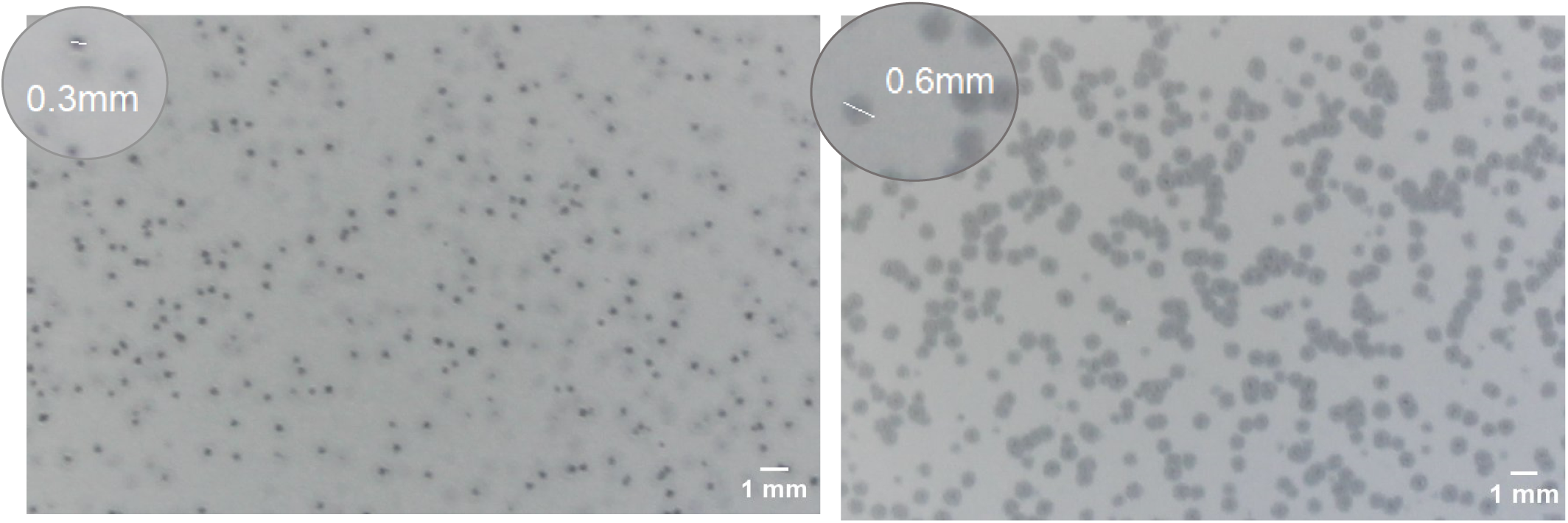
: picture of phage plaques in the layer of soft agar including a sublethal concentration of antibiotic (left, MIC/2) and without antibiotic (right)

### Checkerboard assays

To study the interaction between antibiotic and phage across different concentrations of both agents, we followed a checkerboard assay method as described in previous studies(25,26,39), adapted to our protocol. In a well plate, two-fold dilutions of the antibiotic were distributed in columns and ten-folds dilutions of phage were distributed in lines, before inoculation of the host bacteria. Two methods are applied to evaluate the PAI effect: first, a traditional monitoring of bacteria growth through measurement of OD at 600 nm; second, a measure of bacteria survival by addition of TTC (2,3,5-Triphenyl Tetrazolium Chloride), reduced in formazan red by viable cells, at the end of incubation and measurement of OD one hour after inoculation. We thus have information regarding both kinetic and survival rate at a final point. Since we can see appearance of bacterial resistance starting from the 9^th^ hour of incubation, incubation is stopped and TTC is added at the 8^th^ hour of incubation.

Examples of the kinetic curves are presented in **Error! Reference source not found.**. Full data are available in Table S2. We also present results in the form of an interaction plot which highlights both the effects of the antibiotic and the phage depending of their relative concentrations (Figure 4). This kind of plot has been described by Gu Liu *et al.*(26) to conclude about PAI : both antimicrobials are tested at different level (can be absence or presence) and effect on a value representing bacteria level is reported onto one plot (Figure 4A). If antimicrobials do not impact each other (indifference), the two bacteria levels will follow the same pattern. If there is synergy, bacterial level will be significantly more reduced under combination than under only one of antimicrobial, making the curves on the plot to split apart. If there is antagonism, bacterial level will be enhanced under combination compared to under one of the antimicrobials, making the curves on the plot to cross one another. On our experiments, measured parameter to represent bacterial level is relative area under curve (AUC) (Figure 4B) and relative absorbance (Figure 4C), meaning values for each combination of phage and antibiotic compared to control’s.

**Figure 3.**
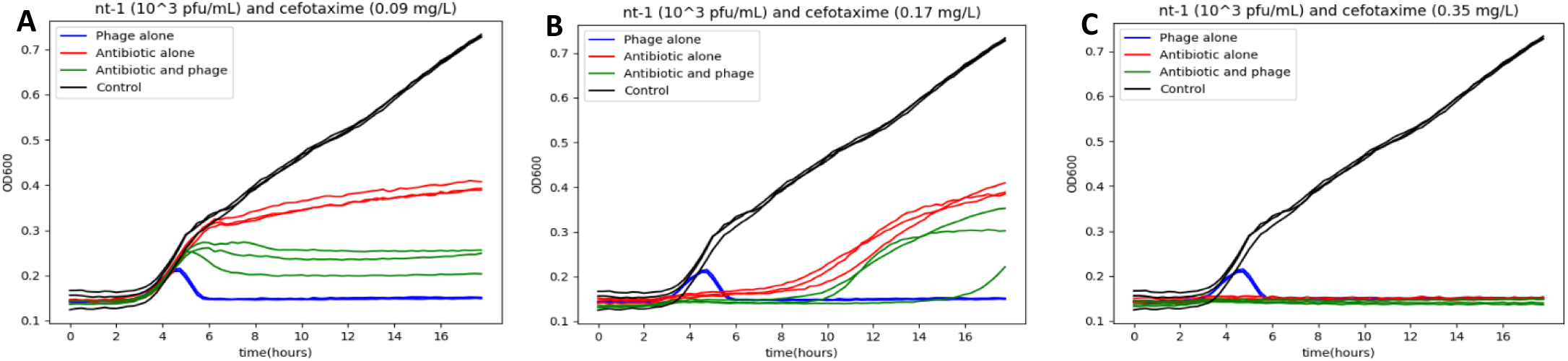
: bacteria OD_600_ kinectic curves, in presence or absence of phage and antibiotic at different concentrations

**Figure 4.**
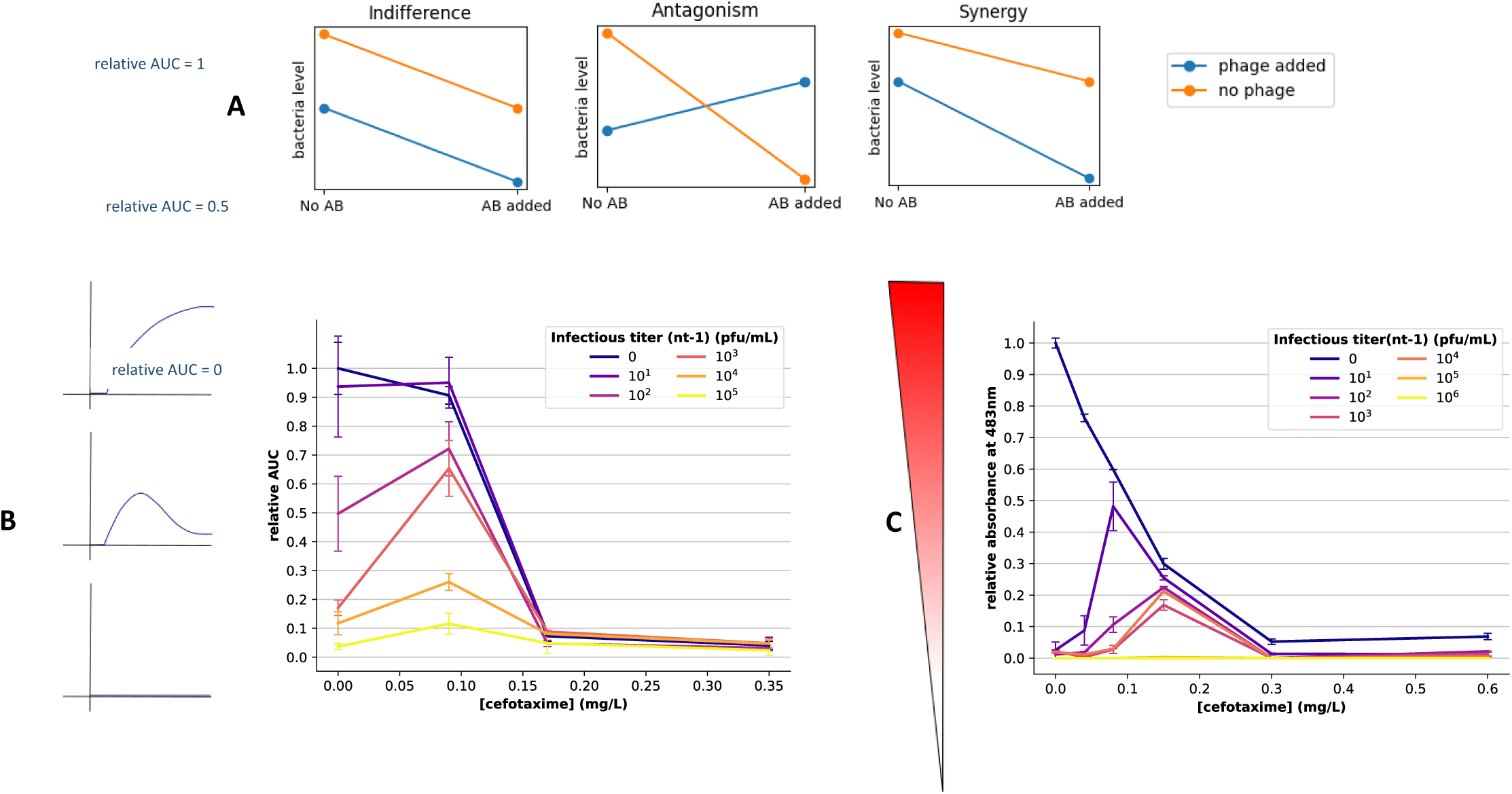
: A) Theoretical interaction plots showing different phage-antibiotic interaction scenarios. B) Experimental interaction plot: normalized AUC of the OD kinetic curves (y-axis) for different concentrations of the antibiotic (x-axis) and phage infectious titres (different curves). Values of triplicates are averaged and divided by the control’s average. Error bars correspond to the standard deviation of triplicate values. C) Experimental Interaction plot: red formazan formation measured from absorbance at 483 nm one hour after TTC (2,3,5-Triphenyl Tetrazolium Chloride) addition, for the different concentrations of the antibiotic and phage infectious titre. Values of triplicates are averaged and divided by control’s average. Error bars correspond to the standard deviation of the triplicates’ values.

From OD_600_ experiment, we observe that inhibition is reached for an antibiotic level of 0.15 mg/L, regardless the presence or absence of the phage. Thanks to the TTC experiment, we can see that despite this absence of growth, bacteria still show evidence of metabolic activity, while this activity is no more detectable from a threshold of 0.3 mg/L of cefotaxime. In the absence of phage for both experiments, the antibiotic presents, as expected, an inhibition effect positively correlated to its concentration, as can be seen by absorbance values decreasing following the x-axis (Figure 4C).

Host and phage are competing in an exponential growth race which phages generally ends up winning. Before bacteria population collapses, kinetic curves shows evidence of an initial growth spurt that is more and more important – in duration as well as in maximum value – as the initial phage titre is lower, a common phenomenon observed in many studies(34,40) (Figure 5). Values of AUC in absence of antibiotic displayed on Figure 4B highlight this phenomenon. Eventual collapse is highlighted on interaction plot (Figure 4C) by all values for no antibiotic and presence of phages being below the relative limit of detection (rLOD) of 0.1 (see material and methods for details on rLOD calculation).

**Figure 5.**
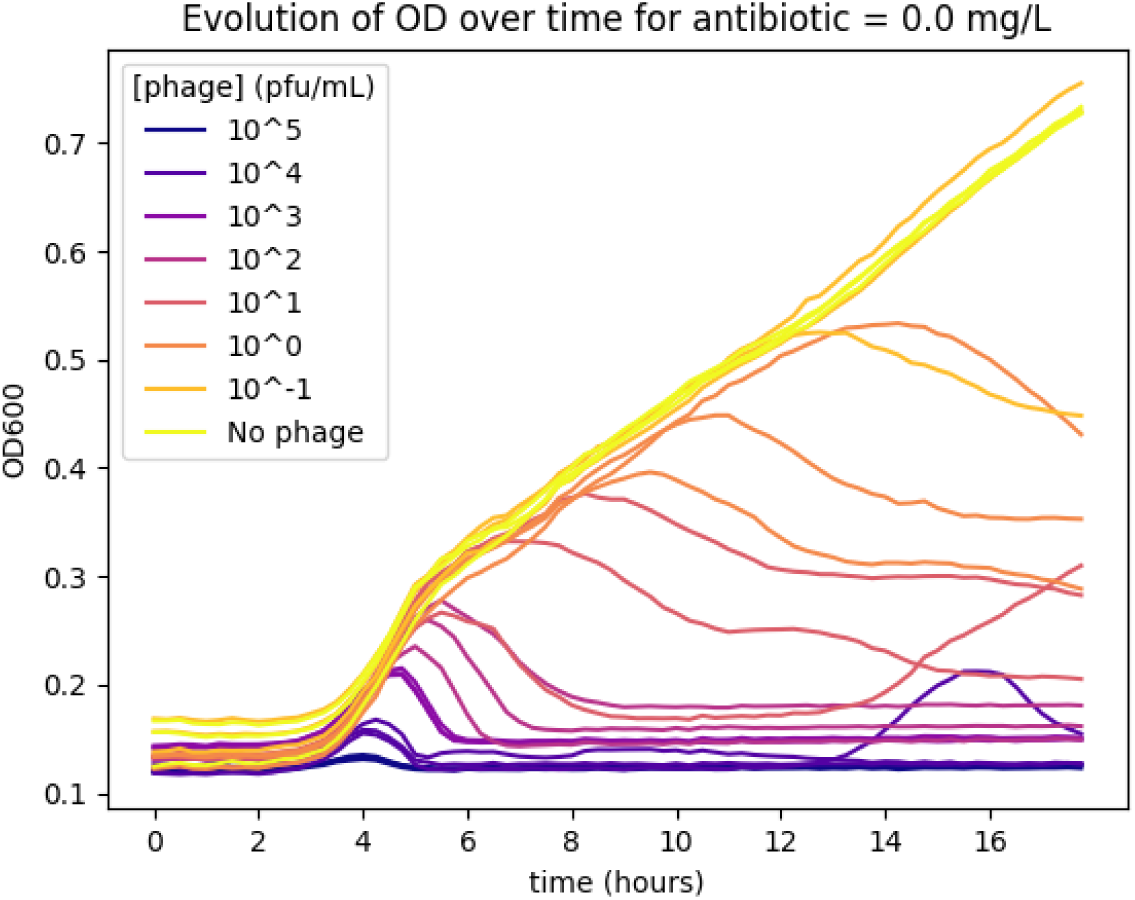
: kinetic curve in presence of various initial phage titres (different curves).

However, at sublethal concentrations of the antibiotic (<0.3 mg/L), the bacteria show evidence of better resistance to the phage (Figure 4B and C). At cefotaxime concentration of 0.09 mg/L, AUC values of combinated curves are higher than in presence of phages alone: this suggests that bacteria elimination is less efficient in the presence of both actors than when administrating the phage alone. On the other side, AUC values of combinated curves are equal or below the ones of antibiotic alone, implying that the phage is not negatively impacting the antibiotic’s action but adding to its effect. At final point, while no metabolic activity is detected in presence of phages, whatever the initial phage titer, survival rate goes up to 10% with addition of antibiotic concentration of 0.15mg/L. Moreover, data suggests that the antibiotic concentration at which this antagonistic effect is maximum may vary depending of the initial phage titre: bacterial survival is even higher (>40%) in wells where initial phage titre of 10 pfu/mL and antibiotic concentration is 0.08 mg/L. Below this optimum, antagonistic effect decreases until being undetectable as initial phage titre increases.

In summary, results suggest that antibiotic at sublethal concentration may have antagonistic effect on the phage’s action, but not the reverse. Antagonistic effect does not appear to be linear but rather undergoes different regimes depending of the antibiotic and phage initial concentration. Yet, we can notice a tendency similar to interaction plots depicted by Gu Liu *et al.*(26): the curve corresponding to the effect of the antibiotic alone is going down as antibiotic concentration increases, while curves representing the effect of combined antibiotic and phage are going up at low antibiotic concentrations, resembling the antagonism interaction scenario (Figure 4A). To further understand this antagonistic effect, we need to get a closer look at phage’s level: in next experiment, we compare virulence factors of phage in presence and absence of sublethal antibiotic concentration.

### One-step growth assay

*To determine the effect of the antibiotic on phage virulence, we carried a one-step growth assay* (*32,33*)*, in absence and in presence of the antibiotic at concentration of 0.17 mg/L (corresponding to half the Minimal Inhibition Concentration (MIC)). An example of results is provided in Figure 6 and the summary of results is provided in from the moment of inoculation of nt-1. The infectious titer is expressed as ×10^6^ pfu/mL*

|  | Latency time (min) | Burst size |
| --- | --- | --- |
| [CTX]=0 | **28 [25;30;30]** | **63 [46;59;86]** |
| [CTX]=0.17 mg/L | **34 [33;34;35]** | **6 [3;6;10]** |

**Figure 6.**
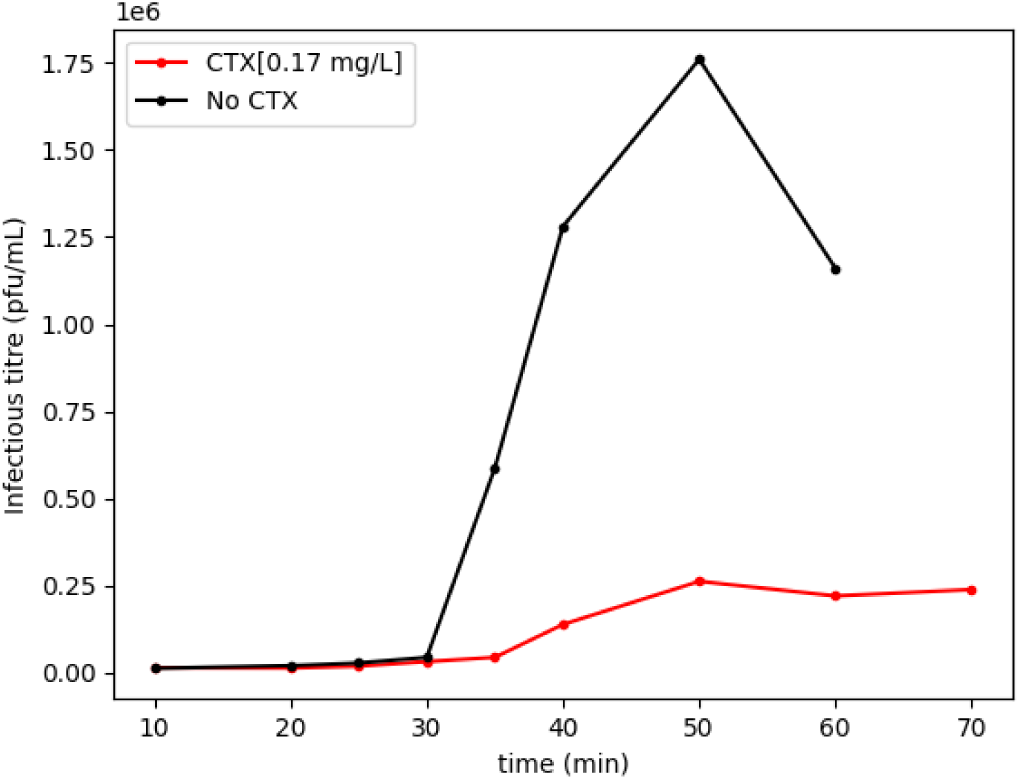
: One-step growth curve in absence (black) and presence (red) of the antibiotic at subinhibitory concentration of 0.17 mg/L from the moment of inoculation of nt-1. The infectious titer is expressed as ×10^6^ pfu/mL

Latency time is slightly delayed in presence of CTX. Although that delay is not that important (6 min), it is close to values found in similar studies from the literature which showed that T4 phage lysis is delayed by 5 min in presence of cefotaxime or ciprofloxacin(36). However, in those studies, the delayed lysis of the T4 coliphage is correlated with an increased burst size, while in our results, the burst size of the vibriophage nt-1 is reduced in presence of the antibiotic by a factor of ten.

## DISCUSSION

### Phage-focused methods

According to Comeau *et al.*, the increased size of lysis plaques in presence of sublethal concentrations of antibiotic is the sign of PAS: antibiotic would increase the burst size of the phage.

In our experiment, we can observe that lysis plaques are smaller in the disk area where bacterial growth is partially, but not fully, inhibited, and larger at the outside border of that zone. It seems reasonable that within the zone where bacterial growth is partially inhibited, phages cannot spread as much as in the control zone: antibiotic, by impeding bacteria growth, also hinders phage growth. As for the larger plaques at the edge of the zone, perhaps a part of the answer lies in the works of Streisinger *et al.*,(41) which shows that the actual area where bacteria are infected might be larger than the visible lysis plaque. The actual extent of phage influence could be revealed by a treatment that weakens the bacterial membrane, such as chloroform vapour, to enable the intracellular phage lysozyme to lyse infected - but not yet lysed - bacteria. In our experiments, there could be a similar effect at stake at the edge of the inhibition zone: cefotaxime at a low concentration would help to lyse unlysed but infected bacteria, leading to bigger lysis plaques, without any link with an increase of the number of produced phage or number of infected bacteria. This effect is coherent with beta-lactamine action, that inhibits membrane synthesis, but would not be observed with other type of antibiotic such as quinolones or aminosides. This hypothesis is supported by the results of gelose experiments conducted with other antibiotics disks. Other beta-lactamases such as penicillins (piperacillin, piperacillin+tazobactam) and other cephalosphorins (ceftazidime) did show a similar interaction effect as cefotaxime (Table S1), to the exception to carbapenems (imipenem, meropenem). One the other hand, fluoroquinones (ciprofloxacine) and aminosides (gentamicine) present no significant effect.

It is important to note that correlating antibiotic concentration with a specific plaque position on the agar is challenging, as antibiotics diffuse over time until they are uniformly distributed across the plate. In contrast, phages are initially homogenously distributed and immediately interact with their bacterial hosts. Plaques spread following the initial infection event. As bacteria grow and their density increases, this further influences plaque dissemination. The dynamic nature of these processes makes it difficult to determine the precise point at which plaque morphology is affected by what amount of antibiotic. Observing variations in lysis plaque size at the periphery of the low-growth zone around the antibiotic disk may suggest that phage virulence factors are altered under certain conditions. However, identifying these conditions requires a robust modelling approach that has yet to be developed. Similarly, it is not possible as for now to directly extrapolate the effects of different sublethal antibiotic concentrations on the bacterial population from this experiment.

To better understand the interaction between phages and antibiotics, further experiments are needed to correlate the nature and intensity of these interactions with specific antibiotic concentrations. We observed that in the presence of a fixed amount of phage and antibiotic at the minimum inhibitory concentration (MIC)/2, lysis plaques were more turbid. However, conducting such experiments on agar across a range of antibiotic concentrations is time-consuming.

One-step growth assays revealed that antibiotic at MIC/2 concentration reduces phage burst size and slightly increases latency time (Figure 6, Table 1). Nevertheless, extending this approach to multiple antibiotic concentrations would be even more labour-intensive.

**Table 1.** : Mean values of latency time and burst size measured in the absence (first row) and presence (second row) of the antibiotic at subinhibitiory level (0.5×MIC). The three repetitions values are presented between brackets.

|  | Latency time (min) | Burst size |
| --- | --- | --- |
| [CTX]=0 | **28 [25;30;30]** | **63 [46;59;86]** |
| [CTX]=0.17 mg/L | **34 [33;34;35]** | **6 [3;6;10]** |

**Table 2 :**
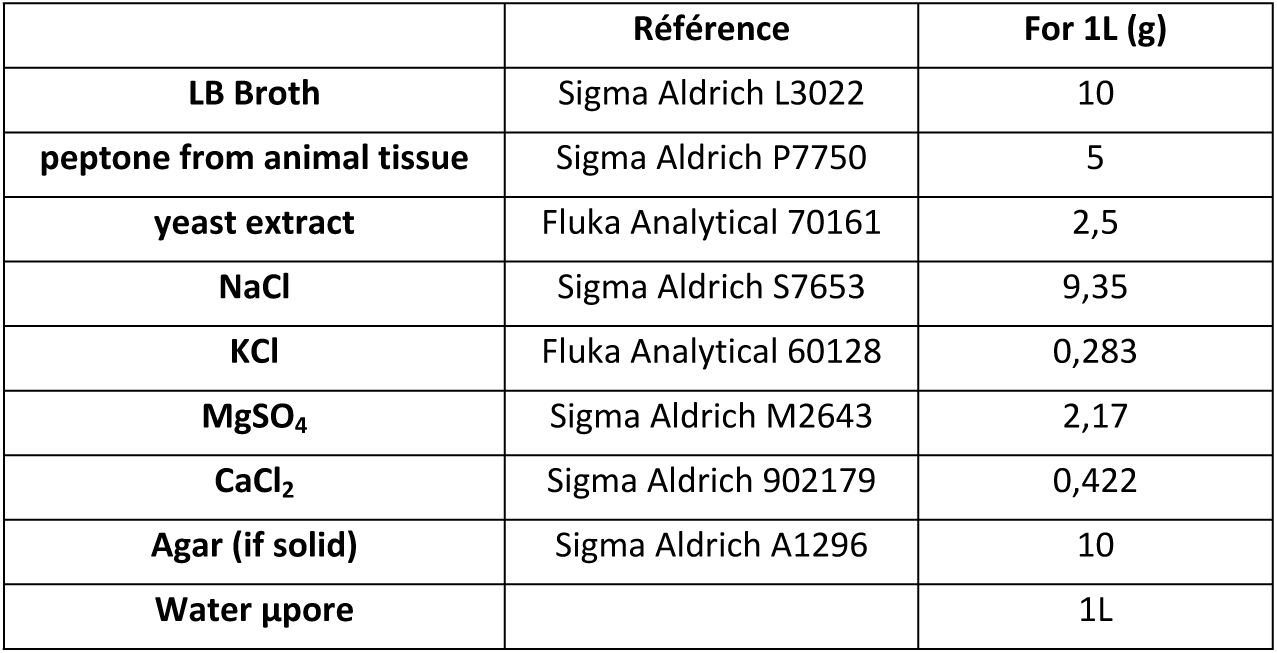
composition of 4-Salt Broth and 4-Salt Agar media.

|  | Référence | For 1L (g) |
| --- | --- | --- |
| **LB Broth** | Sigma Aldrich L3022 | 10 |
| **peptone from animal tissue** | Sigma Aldrich P7750 | 5 |
| **yeast extract** | Fluka Analytical 70161 | 2,5 |
| **NaCl** | Sigma Aldrich S7653 | 9,35 |
| **KCl** | Fluka Analytical 60128 | 0,283 |
| **MgSO<sub>4</sub>** | Sigma Aldrich M2643 | 2,17 |
| **CaCl<sub>2</sub>** | Sigma Aldrich 902179 | 0,422 |
| **Agar (if solid)** | Sigma Aldrich A1296 | 10 |
| **Water µpore** |  | 1L |

To conclusively determine the nature and intensity of phage-antibiotic interactions, biomass-focused methods, particularly those that can be multiplexed, are essential. These methods allow for the assessment of actual effects on the bacterial population, which is crucial for phage therapy, and enable testing across different concentration ranges to identify optimal conditions for maximizing synergy or minimizing antagonism.

### Biomass-focused methods

The most classical way of following bacteria growth, and studying any factor that may influence it, is optical density at 600 nm. This measurement is linearly correlated to the evolution of bacterial biomass, up to a threshold of around 0.1(42). However, measurements are typically taken well above this threshold, and our experiments are no exception.

While measuring the area under the kinetic curve can be a way of summarizing bacterial growth into a singular value, this method shows some limitations. In our case, the AUC value fails to accurately represent the variability in the profiles of the kinetic curves. Antibiotics, whether used alone or combined to phages, lead to delayed growth and lower plateau compared to control (Figure 3). In contrast, phages used alone produce bell-shaped curves, characterized by an initial growth phase followed by a collapse (Figure 5). Indeed, phage growth follow bacteria growth and eventually overcome them. Those discrepancies are hidden when using AUC values alone.

Furthermore, OD is difficult to compare across experiments where objects do not keep given size and shape. Bacteria subjected to sublethal concentrations of antibiotic are unable to properly divide and become filamentous (Figure 8): in such a case, the biomass increase may be due to filamentation rather than cell multiplication. Cell multiplication increases OD linearly (number of diffusing objects), while filamentation lengthens cells, which increases the hydrodynamic radius of bacteria and causes much higher increments of OD (the Rayleigh scattering cross-section varies like the particle radius to the 6^th^-power *r*^6^ (43)). Hence, while looking at combination values, method fails to highlight a comparative effect of each biological agent on the growth dynamic. OD-based metrics such as an inhibition percentage or a FICI value may not be relevant in such a case.

**Figure 7.**
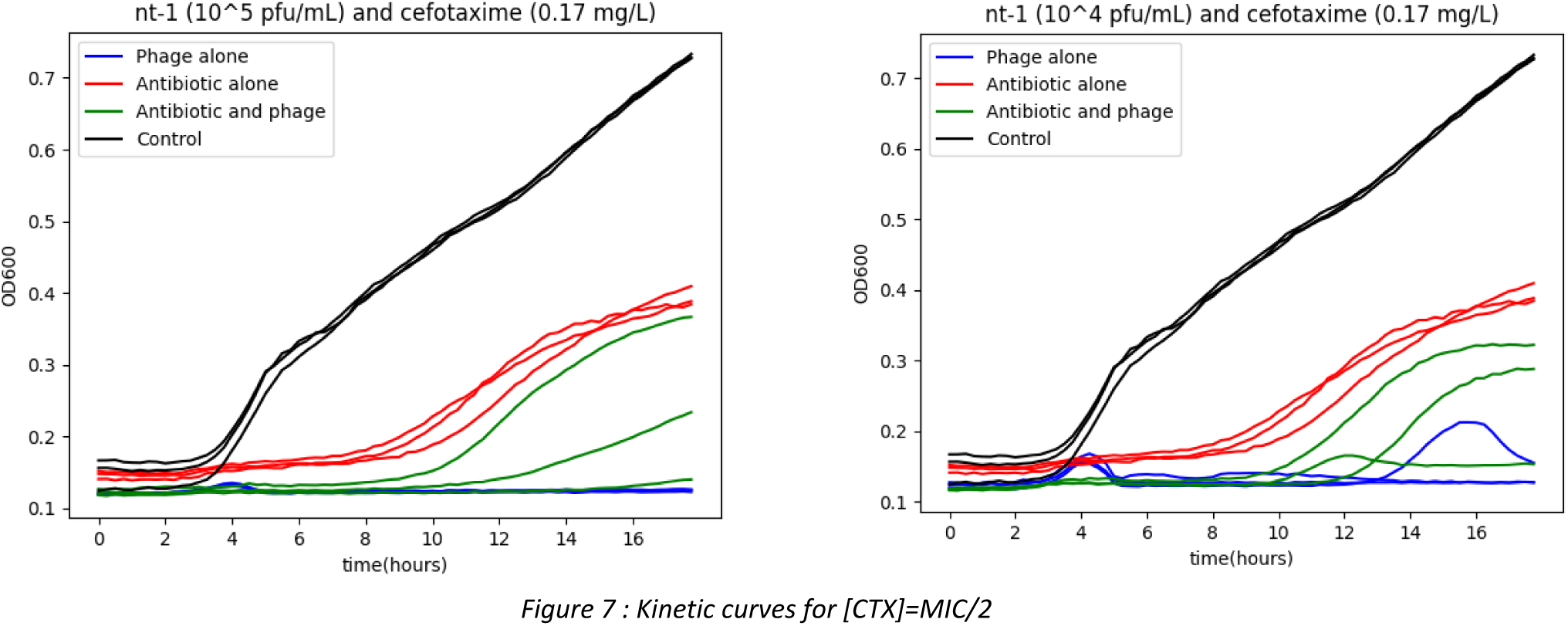
: Kinetic curves for [CTX]=MIC/2

**Figure 8.**
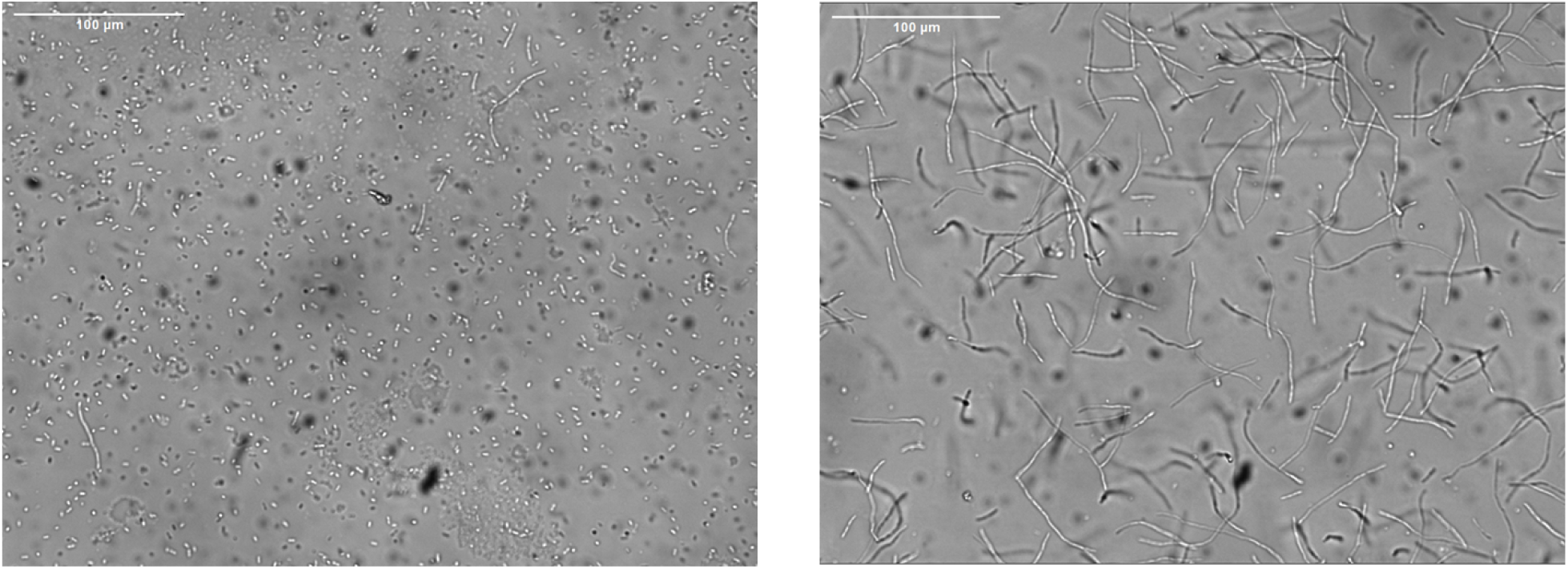
: optical microscopy (x20) observation of bacteria in the absence (left) and in the presence (right) of 170 µg/L of cefotaxime.

It is thus of high interest to support OD_600_ results by the measurement of viability at a final point such as the one obtainable from the colorimetric assay of respiratory activity (red formazan formation). This method allows to obtain a percentage of inhibition from the absorbance measured at 483nm, an optimal wavelength for the quantification of the generated formazan. It is then tempting to summarize interaction into a single value by, for example, computing a **Fractional Inhibitory Concentration Index**(25) :

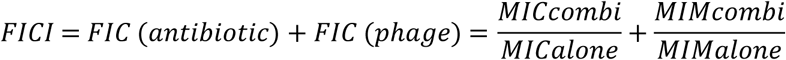

- MIC_alone_ the minimal inhibitory concentration of the antibiotic alone,
- MIC_combi_ the minimal inhibitory concentration of the antibiotic in the presence of phages,
- MIM_alone_ the minimal inhibitory MOI of the phage alone,
- MIM_combi_ the minimal inhibitory MOI of the phage in the presence of antibiotics.

The first term of the equation (FIC(antibiotic)) represents the effect of the phage on the antibiotic, while the second term (FIC(phage)) represents effect of the antibiotic on the phage. Based on this factor value, conclusions can be drawn regarding the type of interaction : synergy (<1), indifference (between 1 and 2) or antagonism (>2)(22).

However, to compute the FICI value, we need to determine at which point bacterial growth is considered actually “inhibited”. One significant hurdle is that phage presence eventually leads bacterial population to collapse, regardless of the phage’s initial titre. Hence, MIC_combi_ cannot be computed since bacteria are inhibited as soon as phages are present whatever their initial titre – it tends to 0. To determine the effect of phage on antibiotic efficacy, survival rates at final point and raw OD_600_ curves are more insightful. The elimination of bacteria is less efficient in presence of the antibiotic alone than with both repressors (Figure 4C), so we can conclude that the phage does not negatively impact antibiotic action but rather allows for an additive elimination effect. However, we can conclude on the effect of the antibiotic on the phage based on the second term of equation. MIM_alone_ value is <10^-4^ (10/10^5^). To determine MIM_combi_, we need to choose at what antibiotic concentration to calculate it, since combinated effect is not linear and seems to have optimum for a certain antibiotic concentration. With cefotaxime at 0.5*MIC = 0.15mg/L, MIM_combi_ goes up to 1 (10^5^/10^5^). The ratio of the latter over the former is >10^4^, which shows strong antagonism of the antibiotic on the phage effect.

It would be interesting to adapt PHORCE model by Mulla *et al.*(34) to link biomass evolution to phage production : however, this model is only applicable if all growth curves present a rise-and-fall behaviour with a clear collapse time, which is not our case.

### Hypothesis on the antagonism of cefotaxime with nt-1

#### Host resistance factors

In all checkerboards assays, we deliberately set the time frame (up to eight hours after inoculation) in order to ignore any bacterial growth appearing after the initial phage-induced collapse, as this behaviour is dependent on the initial presence of resistant strain in the well, a parameter that could vary across repetitions. However, our data suggests that the antagonistic effect of the antibiotic on phages may also appear after eight hours: this late growth spurt seems more likely to appear in the presence of the antibiotic at MIC/2 (Figure 7). This behaviour could mean that the antibiotic, by modifying bacterial membrane, favours the expression of phage resistance factors in its host. In *Enterobacterales* such as *Vibrio*, susceptibility to cephalosporins and carbapenems is often associated with membrane properties(44). Membrane proteins such as outer-membrane proteins (OMPs) are essential to maintain the integrity and selective permeability of the bacterial membrane and play a key role in adaptive responses of Gram-negative bacteria such as solute and ion uptake, iron acquisition, antimicrobial resistance, serum resistance. As a consequence, the expression of OMPs depends on antibiotic stress and it is known at the same time that some of them are the receptor for vibriophages(45–47), as well as for other Enterobacterales(48). Further investigation of this hypothesis would require to decide the delay after which to stop the measurements, a difficult task as we do not know how late resistant strains can start to grow.

#### Uneven production within bacteria population

Cefotaxime causes the bacteria to grow filamentous (Figure 8). In our experiments, as a consequence of that filamentation, the antibiotic acts as antagonist toward the phage. The strength of the antagonism effect varies depending of the antibiotic concentration. Lysis plaques can appear bigger or on the contrary, more turbid depending on the concentration and diffusion of the antibiotic in agar. In any case, it appears that the antibiotic is able to favour host bacteria resistance to the phage in liquid medium by decreasing drastically the phage’s production rate.

It is noteworthy that this bacterial decrease is observed at the population scale and may not reflect individual behaviour. Indeed, such a decline could either stem from an inhibited productivity of each individual bacteria, or from a lower number of infected cells that actually produce phages. Based on the lysis pathway, we propose the following hypotheses regarding the antagonism mechanism: filamentation in some bacterial cells may impede holin accumulation to the critical threshold required for endolysins to degrade the cell membrane. Consequently, lysis may be delayed or even prevented in a subset of the population, while lysed cells may exhibit a burst size similar to or even greater than that observed under control conditions. This bimodal distribution within the population could result in an overall reduction in productivity.

However, it is challenging to assess the robustness of this hypothesis, as the critical threshold of holin required for lysis remains unknown. Most studies on the interplay between holin, antiholin, and endolysins have focused on the lambda phage, and quantitative estimates of the involved components are yet to be determined. Further research, potentially involving mathematical modelling, is needed to elucidate the dynamics of these interactions and validate the proposed hypothesis.

#### Lysis inhibition

Another explanation could be based on the hypothesis that the phage is able to delay its own lysis under some conditions, a phenomenon known as lysis inhibition (LIN). This phenomenon is observed in response to secondary adsorptions: latent period of the phage is extended, leading to higher burst when host is eventually lysed, since progenies had more time to mature(41,49). Such a phenomenon has been mainly studied in T-even coliphages, but Hays *et al.*(50) proved that some vibriophages are also able of such a behaviour. We can presume secondary infections are more likely to happen on filamentous bacteria. Turbid plaques, as the ones we can observe in antibiotic-enriched agar, can also be the evidence of LIN(41,51). LIN is usually expected to resolve into a later higher burst, providing a significant advantage over strains not able of such a behavior(41,51). However, in our case, that later burst never happened: we sampled from one-step growth assay for 24 hours without reaching a higher value on numeration plates than the one obtained at 60 min (Figure S3). That does not invalidate LIN hypothesis, though: it has been observed that delayed lysis might lead to no lysis at all(51). That can happen if superinfection keep occurring within <10 min intervals(52), when nutrients run out(51), under phage satellite action(50) or because of genetic mutation(53). In our case, starvation upon exhaustion of nutrients seem the most probable event that could have trigger infinite lysis inhibition.

Overall, it seems that the nature of PAI for a given phage/host/antibiotic cannot be reduced to a one-word conclusion such as “synergy” or “antagonism” and/or one FICI value. The nature and intensity of interaction highly depends on many factors, antibiotic concentration being the most prominent. Our *in vitro* results in liquid medium suggest that there exists an optimum concentration of antibiotic that induce maximum phage-antibiotic interaction, in our case, antagonism between cefotaxime and nt-1 phage. Another significant factor that might affect these observations is the timing of phage administration relative to antibiotic exposure. In our liquid experiments, both the phage and antibiotic were administered simultaneously. If the phage was introduced prior to the antibiotic, allowing a window of, for example, two hours, the observed antagonistic interactions might be reduced or altered. This aspect warrants further investigation in future studies to determine the optimal sequencing of treatment for maximum efficacy.

Agar experiments serve as an accurate, cost-effective, and straightforward initial approach to observe potential interactions. However, they are too limited to easily correlate observed plaque morphology with quantitative PAI effects. To extract more information from agar experiments, several solutions can be proposed:

- One approach to obtain more detailed information is to follow up the experiment with chloroform vapor treatment(41). The potentially extended lysis plaques resulting from this treatment can provide insights into the actual diffusion of phage infection.
- The E-test(27), in comparison with standard agar experiments, could provide a more quantitative solution and is worth further investigation. It could help determine if the minimum inhibitory concentration (MIC) of an antibiotic is affected by the presence of phages, or vice versa.
- Lastly, we can imagine developing a supervised learning model. This could involve conducting a series of experiments on agar, using antibiotic disks on plates seeded with phage and bacterial hosts, along with liquid-based experiments using a checkerboard method. The goal would be to predict the outcomes of the liquid-based experiments based on the results from the agar experiments. This strategy leverages the advantages of agar experiments, which are easy, cost-effective, and scalable, to draw robust conclusions about phage-antibiotic interactions (PAI).
To get a broader overview of PAI, it is essential to employ multiplex testing methods such as checkerboard assays. Additionally, combining techniques like kinetic monitoring and endpoint viability measurements can provide more detailed insights. Traditional assays in well plate can test up to 96 different conditions, which is pretty low comparing to automats such as Vitek2, that can test up to 18 different antibiotics with a range of 4 different concentrations. The future of phagotherapy belongs to those able to widely automatize PAI tests and use highly multiplexing methods such as droplet-based millifluidic systems(54).

## MATERIALS AND METHODS

### Medium and strains

*Vibrio natriegens* of ATCC® 14048™ strain from American Type Culture Collection were used in this study. The nt-1 phage was bought from Centre de référence pour les virus bactériens Félix d’Hérelle, Laval, Canada. Media of infection were 4-Salt Broth for liquid phase and 4-Salt Agar for solid phase (Table). Phages were amplified in liquid medium and purified by centrifugation and filtration at 0.2 µm as described previously. Infectious titres were measured by inoculating phage solutions diluted in NaCl 0.9% along with overnight host cultures in 4-Salt agar plates and counting the number of lysis plaques after 18 hours of incubation at 30°C. Phage suspensions in NaCl 0.9% were stored at 4°C.

### Experiments in agar

A few colonies of bacteria were picked up from agar plates and suspended in 4-Salt broth medium to grow overday at 30°C under agitation until bacterial concentration reached 10^9^ cfu/mL. 200 µL of that solution were inoculated along with 100 µL of the phage at 10^5^ pfu/mL in 4-Salt agar medium and poured on an empty dish. Cefotaxime, diluted in Millipore water to reach the desired concentration, was added either directly in agar or on a 6-mm disk (FILTRATECH PA320A0006). Plates were left to dry, incubated at 30°C and read after 18 hours of incubation.

### Image processing of lysis plaques on agar

The image of the Petri dish was a 3300 × 3300 pixels 8-bit (255 gray levels) image. It was taken by CANON EOS600D camera, with 18-55mm objective. Image processing was implemented with Python and the OpenCV library. The consecutive steps to obtain the density of lysis plaques as a function of the distance to the antibiotic disk were the following:

- Pre-processing: A high-pass filter was applied to the images via the convolution with a Gaussian kernel with σ
= 3 pixels. This step allowed to eliminate most of non-significant defects of the agar surface.
- Thresholding: The image was then thresholded to a binary image, *i.e.* all pixels with a gray value above a given threshold were considered to represent lysis plaques. As the gray level of the agar was heterogeneous across the plate due to the varying bacterial concentration, we implemented local thresholding to account for these discrepancies, meaning that the threshold was not set globally for the whole image but rather locally in sliding windows. This local thresholding method used a window size of 201 pixels (corresponding to around 0.5% of the Petri dish area), and the resulting local threshold was set to be the mean of that window (ponderated with a Gaussian kernel) minus 8 gray levels.
- Computing radial density: Circle selections of increasing diameter, centered on the antibiotic disk, were considered. For each selection, the lysis plaque density was computed as the ratio between thresholded pixels (i.e., those corresponding to lysis plaques) and the total number of pixels on the circle.
- Post-processing results: The 1-D array of density values as a function of distance to antibiotic disk was smoothed using a Savitzky-Golay filter with window size 51 and polynomial order 3.

### Checkerboard assay: area under curve (AUC) of the OD_600_ kinetic curve

#### Plate preparation

In each well, 10 µL of 10-fold dilutions of phages from 10^-1^ to 10^5^ pfu/mL were distributed among rows. 10 µL of 2-folds dilutions of antibiotics from 0.09 mg/L to 0.35 mg/L were distributed among columns in triplicates. One column was dedicated to phage controls (10 µL of NaCl 0,1% instead of phage) and three columns to antibiotic controls (10 µL of pure sterile water instead of antibiotic). Three wells contained bacteria without any antibacterial agent (no phage, no antibiotic). See supplementary Figure S1 for full schema of the well plate. 20 µL of host bacteria was added to each well at concentration of 2*10^5^ cfu/mL for a total volume of 200 µL in each well. OD was measured at 600 nm every 15 min during 18 hours with a Tecan Infinite M1000 spectrophotometer.

#### Data treatment

Values of OD_600_ at t_0_ were subtracted from values of OD_600_ at each time point in each well to compensate for variations in the initial inoculums. From those adjusted values, the area under kinetic curves (AUC) are measured for each well. Since we can see appearance of bacterial resistance starting from the 9^th^ hour of incubation, AUC is measured up to the 8^th^ hour of growth. Values of AUC for each well normalized by the AUC of controls (*i.e.,* without either phage or antibiotic) are calculated as following:

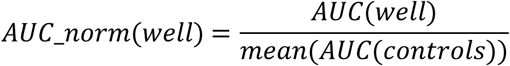

AUC(control) is the averaged AUC of control wells (no antibiotic, no phage). Detailed data are available in supplementary. Values of AUC_norm are averaged for triplicates and reported on interaction plots for each antibiotic and phage concentration.

### Checkerboard assay: OD_483_ after addition of tetrazolium salts

#### Plate preparation

In each well, 10µL of 10 folds dilutions of phages from 10 to 10^6^ pfu/mL were distributed among rows. 10 µL of 2-folds dilutions of antibiotics from 0.04 to 0.6 were distributed among columns in duplicates. One line was dedicated to phage controls (10 µL of NaCl 0,1%) and two columns to antibiotic controls (10 µL of pure sterile water instead of antibiotic). Two wells contained the bacteria without any repressor, with addition of NaCl 0.1% and sterile water instead of phage and antibiotic. Three wells were inoculated with 180 µL medium and 20 µL bacteria as global growth controls. See supplementary Figure S2 for full schema of the plate. Since they eventually absorbed the same as the former two after addition of 2,3,5-triphenyl tetrazolium chloride (TTC), their results were averaged to compute DO_483_ ratio denominators. Three wells were dedicated to phage sterility controls and three others to antibiotic sterility control (no bacteria). Three wells were pure medium sterility control. 20 µL of host bacteria was added to the relevant wells at concentration of 2×10^5^ cfu/mL for a total volume of 200 µL in each well. 20 µL of TTC at 0,1% was added to each well at the 8^th^ hour of incubation. Optical density was measured at 483 nm after one hour.

#### Data treatment

The ratio of OD_483_ was calculated for each well as following:

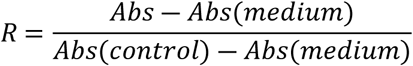

Abs is the absorbance in the well at the wavelength 483 nm.

Abs(medium) is the average of the absorbance of wells H4 to H12

Abs(control) is the average of the absorbance of wells G11, G12, H1, H2 and H3

Duplicates values are averaged and reported on the interaction plot for the different antibiotic and phage concentrations. Full data are available in supplementary.

#### rLOD measurement

The Limit of detection of rLOD measurement is computed as recommended by the American Chemical Society(38)using the following formula:

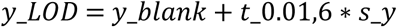

With:

- y_LOD the absorbance value of the limit of detection
- y_blank the average absorbance value of the 7 controls (medium without bacteria)
- t_0.01,6 the Student quantile for 7 repetitions and a significance level α of 0.01, being 3.143
- s_y the standard deviation of the measurements

rLOD, the relative absorbance of the limit of detection, is then measured using same formula than described in Data treatement.

### One-step growth assay

Bacteria were grown in 4-Salt Broth with cefotaxime (CTX) at 0.17 mg/L (half of the minimum inhibitory concentration) five hours before the experiment until it reached an OD_600_ at 600 nm higher than 0.3. The solution was then diluted with the same medium enriched with antibiotic in a 50 mL Falcon tube (adsorption tube) to reach an OD_600_ value of 0.3. The phage solution was added to the adsorption tube at a concentration of 10^6^ pfu/mL. After a five-minute delay at 30°C under agitation in order to let phage infection begin, two other tubes were prepared by dilution of the adsorption tube with dilution factors 10^2^, 10^3^ and 10^4^. Samples were collected from the different tubes at regular time intervals for plaque counting in soft agar to monitor the evolution of the phage infectious titre after adsorption. Burst size was calculated as the ratio between the average value of the first plateau and the average value of the second plateau. Latency time was defined at the time of the intersection between the first plateau and slope curve.

## SUPPORTING INFORMATION

**S1 Table. Phage-antibiotic interaction on agar for different antibiotics.** Soft agar is homogenously seeded with both bacteria and phages before depositing antibiotic disk at the center. The proportion of surface occupied by lysis plaques is then computed as a function of the distance to center.

(DOCX)

**S2 Figure. Checkerboard assay using a 96-well plate for AUC calculation.**

(DOCX)

**S3 Figure. Checkerboard assay for experiments with TTC (2,3,5-Triphenyl Tetrazolium Chloride).**

(DOCX)

**S4 Figure. OSGA (one-step growth assay) with sublethal cefotaxime concentration up to 20 hours.**

(DOCX)

**S5 Table. Kinetic curves of *Vibrio natriegens* in presence, absence and combinated presence of cefotaxime and nt-1 phage.**

(DOCX)

## ACKNOWLEDGEMENTS

We thank Hugues de Villiers de la Noue for his advices and support on the technical aspects.

## DATA AVAILABILITY STATEMENT

The data that support the findings of this study are available from the corresponding author upon reasonable request. The script used for the lysis plaque morphology analysis (Figure 1) is available with the following DOI: 10.5281/zenodo.19736277

## AUTHOR CONTRIBUTIONS

**Conceptualization:** Brunel Archambeaud, Pierre R. Marcoux.

**Data Curation:** Brunel Archambeaud, Clément Douarre.

**Formal Analysis:** Brunel Archambeaud, Clément Douarre.

**Funding Acquisition:** Pierre R. Marcoux.

**Investigation:** Brunel Archambeaud.

**Methodology:** Brunel Archambeaud, Clément Douarre.

**Project Administration:** Brunel Archambeaud, Pierre R. Marcoux.

**Resources:** Brunel Archambeaud, Pierre R. Marcoux.

**Software:** Clément Douarre.

**Supervision:** Pierre R. Marcoux.

**Validation:** Brunel Archambeaud.

**Visualization:** Brunel Archambeaud, Clément Douarre.

**Writing – Original Draft Preparation:** Brunel Archambeaud.

**Writing – Review & Editing:** Brunel Archambeaud, Clément Douarre, Pierre R. Marcoux.

